# Chromatin packing domains persist after RAD21 depletion in 3D

**DOI:** 10.1101/2024.03.02.582972

**Authors:** Wing Shun Li, Lucas M Carter, Luay Matthew Almassalha, Emily M. Pujadas-Liwag, Tiffany Kuo, Kyle L MacQuarrie, Marcelo Carignano, Vinayak Dravid, Masato T. Kanemaki, Igal Szleifer, Vadim Backman

## Abstract

Understanding chromatin organization requires integrating measurements of genome connectivity and physical structure. Prior work demonstrates that RAD21 depletion results in the complete loss of topologically associated and loop domains on Hi-C, but the corresponding change in physical structure has not been studied using electron microscopy. Pairing chromatin scanning transmission electron tomography with Hi-C, we study the role of cohesin in regulating the spatially resolved, conformationally defined chromatin packing domains. We find that only 20% of packing domains are lost on electron microscopy upon RAD21 depletion with the effect primarily on small, poorly packed (nascent) domains. Overall, this contrasts with the prevailing understanding of genome regulation, indicating that while cohesin influences domain formation, non-cohesin mediated mechanisms predominantly regulate the 3D genomic physical structure.

## Introduction

Our understanding of the mechanisms that constrain chromatin organization in 4D space is crucial to resolving how DNA replication, repair, and RNA transcription are regulated(*1–6*). Further, distortion in chromatin organization is associated with multiple disease processes, including numerous malignancies(*1, 2, 5, 7–10*). Since the emergence of high-throughput chromatin conformation capture (Hi-C) sequencing, researchers have worked to understand the mechanisms that result in the formation of topologically associated domains (TADs) and how these domains correspond to physical structures within individual cells(*11–17*). From their initial description as foci of higher-than expected contact probability in ensemble population measurements, several key features of TADs have been described. TADs are predominantly associated with cohesin-mediated loop extrusion in association with CTCF binding motifs and potentially act as boundary mediating elements in supra-nucleosomal chromatin organization(*13, 15, 18–22*). Cohesin loops at individual loci appear to be transient in nature and relatively rare events, residing in a loop conformation <10% of the time in a single locus imaged in mouse embryonic stem cells(*23*). Finally, TAD boundaries appear to be crucial elements in collective genome function as their targeted knock-out results in pathogenic phenotypes during mouse development(*24*). As TADs arise in population measurements, it was proposed that TADs are an ensemble feature of cellular population measurements; as such, they may represent infrequent, but crucial structures necessary for proper collective function(*17, 23*). In support of this, TAD-like structures have been observed with variable positions and frequencies utilizing oligo-paint based super-resolution microscopy (*17, 20–22*) and loss of RAD21 did not disrupt heterochromatin cores on structured illumination(*25*). Despite the association of TADs with cohesin-mediated loop extrusion, RAD21 depletion did not result in the loss of all the TAD-like structures in individual cells (*14, 17, 20–22*). Further, the boundary strengths of TAD-like structures were comparable in the RAD21 depleted cells to those with intact cohesin(*20, 22*). Although oligo-PAINT based super-resolution imaging provides sequence-specific loci information, the resolution limit of this modality approaches ∼30nm and can only target specific loci for study(*14, 17, 20–22*). Further, these methods rely on the formamide-based DNA denaturation, which has been shown to alter nanoscale 3D genome structure on electron microscopy(*26*). As such, it cannot provide insights on the transition of chromatin from the disordered polymer structures observed via chromatin electron tomography (ChromEMT) into higher order structures. Likewise, it is insufficient for providing quantitative information about the global features of chromatin 3D folding throughout the nucleus(*27–29*).

In their seminal work, Ou et al demonstrated that by using photo-oxidation of a DNA-specific dye, osmium tetroxide could be specifically localized to DNA in a density dependent manner(*27*). With this approach, ChromEMT was able to overcome the limitation from prior EM studies of chromatin and allow the direct resolution of DNA, nucleosomes, and fibers. In ChromEM imaging, mass density is proportional to intensity. As a result, unlike super-resolution imaging techniques with comparable resolution, ChromEM resolves the ground-truth structure of chromatin(*27*). Combining ChromEM with scanning transmission chromatin electron microscopy (ChromSTEM) with high-angle annular dark-field (HAADF) tomography, we previously demonstrated that we can thus resolve chromatin structure with a resolution of 2nm and allow direct quantitative analysis of chromatin density across all the regulatory length-scales of the genome in 3D(*28, 29*). Using ChromSTEM-HAADF tomography, we demonstrated that chromatin organization transitions from the disordered polymer structure observed in Ou et al into higher-order packing domains that are 50-200nm in size and composed of ∼100-500kb of DNA; sizes which were remarkably similar to those predicted for TADs and loops(*28, 29*). Whether packing domains form primarily due to loop extrusion processes and how topological associated domains relate to ChromEM-resolved higher-order chromatin structures evades our understanding.

In this work, we perform ChromSTEM tomography on RAD21 depleted cells using the HCT116 RAD21-mAID-Clover CMV-osTIR1(F74G) cell line and pair it with ensemble Micro-C to demonstrate that packing domains are not solely formed due to cohesin-mediated loop extrusion(*30*). Although approximately 20% of packing domains are lost in RAD21 depletion, the remaining domains retain similar sizes and polymeric organization to those observed in the controls, indicating that the majority of genome physical structure form independent of cohesin-loop extrusion. Finally, we observed that functionally, RAD21 depletion is primarily associated with domain assembly, as nascent domains (small, lower density packing domains) are predominantly lost in ChromSTEM tomography. In contrast, mature packing domains appeared largely unaffected by the depletion of RAD21, indicating that a second mechanism of domain formation exists and that, once formed, domains are potentially maintained by other processes. Consequently, it appears that TADs and loop domains represent features of connectivity and may not directly manifest the space-filling structure that arise in chromatin *in vitro*. In summary, this work demonstrates that (1) packing domains are the predominant higher-order structures within the eukaryotic nucleus, (2) packing-domains do not depend exclusively on cohesin mediated loop extrusion to form and function, and (3) topological associated domains likely do not directly map to physical domains.

## Results

### Chromatin organizes into three distinct regimes in colonic HCT-116 cells

ChromSTEM has previously demonstrated that chromatin organization exists across three-hierarchies in A549 pulmonary epithelial cells and in BJ-fibroblast cells: (1) a disordered chromatin polymer (5-25nm), a power-law polymer (50-150nm), and space-filling territorial polymer (>200nm)(*28, 29*). With respect to the disordered polymer, these observations were consistent with the structures previously identified in Ou et al where chromatin organizes as a flexible fiber with variations in density and folding(*27*). Interestingly, this disordered polymer produces a power-law polymeric regime at higher length-scales due to the resulting formation of packing domains. Finally, these structures converge into a territorial polymer with a random spatial arrangement of densities. Notably, this last regime is not an assembly of the underlying regimes but is a random distribution of mass-density(*28, 29*).

To investigate if these regimes further extend in human cells to HCT-116 colonocytes, a model of microsatellite unstable colorectal cancer, we performed ChromSTEM-HAADF tomography utilizing the protocol previously described(*28, 29*). In brief, a major prior limitation of electron microscopy within the nucleus was the result of non-specificity of negative staining agents to both chromatin and non-chromatin molecules within the nucleus. To overcome this limitation, Ou et al demonstrated that utilization of the DNA-specific dye, DRAQ5, with photo-oxidation results in the preferential binding of osmium of DNA(*27*). DRAQ5 stained HCT-116 control cells were resin embedded and cell nuclei were identified utilizing wide-field optical microscopy. We subsequently sectioned 120nm resin section and performed dual-tilt STEM with high annular high-angle annular dark-field imaging on a 1.9um x1.9um section producing a high-resolution tomogram from within the center of the nucleus (**Fig. 1a&b**). As expected, ChromSTEM tomography resulted in the reconstruction of high-resolution features of chromatin including 3-D rendered individual fiber loops (**Fig. 1c, Movie S1**) and chromatin packing domains (**Fig. 1d, Movie S2**). Consistent with prior studies in A549 and BJ-fibroblast cells, chromatin organized into three regimes within HCT-116 control cells with a power-law like geometry within supra-nucleosomal length-scales apparent (**Fig. 1e**) within individual domains.

**Fig. 1.**
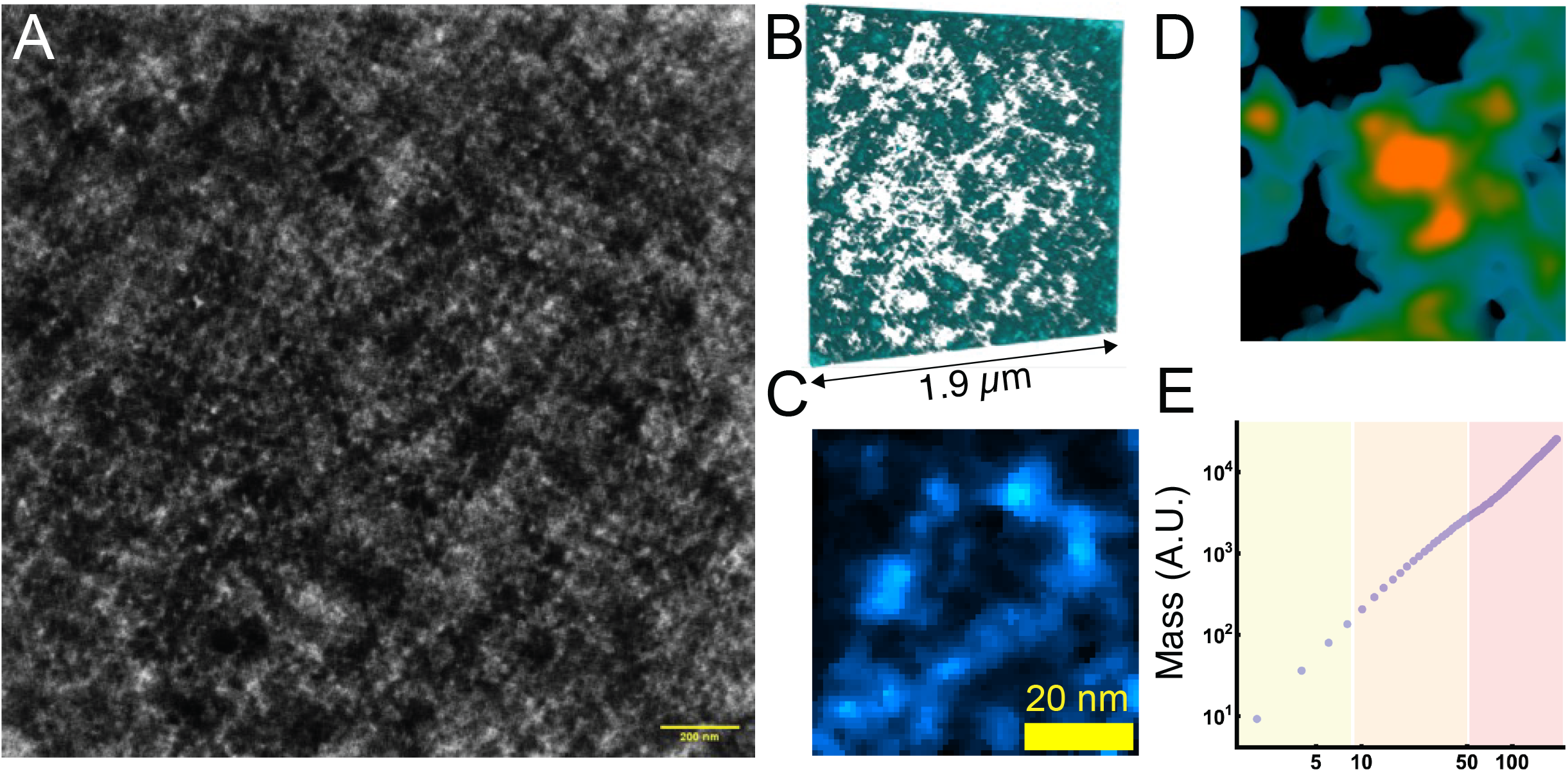
ChromSTEM preparation and tomogram analysis. **(A)** High-resolution mean projection from ChromSTEM in HCT-116 cells. **(B)** Tomogram reconstruction showing distribution of high-density areas with surrounding porosity in 100nm section. **(C)** Visualized chromatin loop with approximate length of 120nm. **(D)** High resolution of packing domain tomogram projection (200nm x 200nm) showing high density within chromatin center with progressively decreasing intensity until porous regions are encountered (black). **(E)** Log-log plot of mass density distribution vs. radius from the visualized packing domain demonstrating the emergence of three distinct chromatin regimens: Yellow (disorder-polymer), Orange (power-law polymer), Red (territorial polymer). The transition between states occurs throughout the nucleus and varies between packing domains.

### Packing domains are heterogeneous, higher order supra-nucleosome structures

Utilizing the produced ChromSTEM tomogram, we subsequently performed analysis to evaluate the organization of structure into packing domains. In prior work, it was shown that packing domains are heterogeneous structures with a distribution of sizes, densities, and packing efficiency. A characteristic feature of packing domain organization is that the density distribution follows a power-law geometry characteristic to polymeric structures such as chromatin that is detailed below. To identify packing domains, we first analyze the mean-projection of the ChromSTEM tomogram to identify the loci with the highest local density as defined by the 1.5-times the standard deviation of the local density. The size of packing domains is then defined by the smallest radius from three independent classification measurements: the radius where deviation of density from the log-log density-size distribution occurs (i.e., where it no-longer follows power-law mass density behavior), the radius at which the first derivative reaches a space-filling geometry, and the radius at which the minimum of density occurs (**Fig. 2a, Fig. S1**). In this approach, packing domains (pink circles) with their local centers (white points) are identified for further analysis (**Fig. 2a**).

**Fig. 2.**
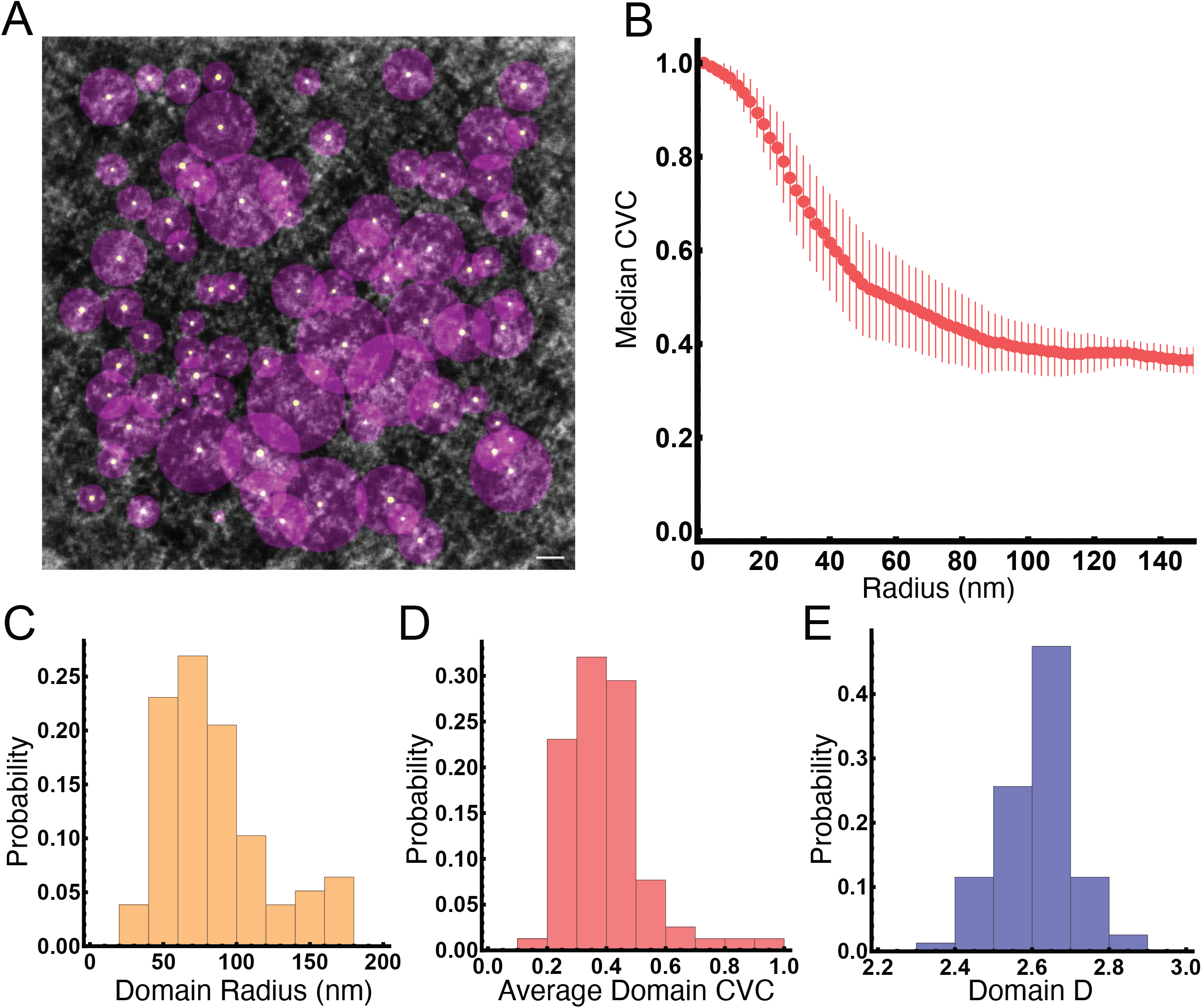
Chromatin organizes into packing domains in HCT-116. **(A)** Packing domains identified in ChromSTEM tomogram with projection of their centroid (white circle) and bounding radius (purple) in control cells. In total, 78 packing domains were identified within this tomogram. Packing domains are identified by local threshold detection and the radius is determined by the minima of either the (1) radius deviates from log-log density, (2) radius that reaches the first derivative or (3) the radius at which the minimum density occurs. **(B)** Chromatin volume fraction decays from the center of packing domains toward their periphery, with the PD density approaching the average CVC of the nucleus near the periphery of the packing domains. Errorbars represent median deviation. **(C-E)** Packing domains are heterogeneous structures with a distribution of sizes **(C)**, CVC **(D)**, and power-law packing **(E)**.

We observe that packing domains demonstrate the highest density within the center (chromatin volume concentration (CVC) ∼1 at the interior core) with the subsequent decay to the nuclear wide average CVC of ∼0.3 (**Fig. 2b**). As expected, PDs have a distribution of sizes (**Fig. 2c**, average radius of 84nm with standard deviation of 36nm) and average chromatin volume fractions (**Fig. 2d**, average CVC of 0.40 with standard deviation of 0.13) As a polymeric structure, the relationship between the mass density distribution of chromatin and its shape can be quantified either by the contact scaling relationship, *S*, or in relation to the mass-distance relationship with fractal dimension, *D*, as follows. *S* measures the frequency of contacts between ‘monomers’ in relation to the linear distance on the chain and decays as a function of distance depending on solvent conditions, confinement, crowding, and other considerations. *D* is a complementary measure of the polymer which relates the distribution of mass to the occupied volume as a function of the radial distance, *r*, by *M* ∝ *r*^*D*^. As with *S*, the measured *D* for a chromatin polymer depends on the solvent conditions, monomer-monomer interactions, crowding, etc. It is well established that chromatin is not in the limiting case of a collapsed, space filling globule structure where *D*=3 nor is it a polymer in idealized solvent conditions where the monomers prefer interactions with the solvent where *D*=5/3. Instead, as has been observed in multiple experimental modalities, chromatin behaves as a power-law polymer where interactions with the monomers are preferred resulting in *D* ranging between 2 and 3 at supra-nucleosomal length-scales.

Utilizing the relationship between mass and distance, we calculated the fractal dimension, *D* within packing domains and found that, as expected, they deviate from a purely space-filling geometry with an average *D=* 2.61 (standard deviation of 0.09, **Fig. 2E**) in control HCT116-RAD21-AID2. In conjunction with prior studies, this suggests that supra-nucleosomal chromatin organization is not simply a polymer in a good solvent (D=5/3), nor does it resemble a purely-space filling globule (D=3) but is instead a disordered polymer assembly with a maximum observed *D* of 2.84 (**Fig. 2E**). A notable feature of packing domain structure is the corrugation, indicating that the continuous distribution of chromatin transitions from chromatin dense regions to porous open segments (**Fig. 1d**) that could be accessible to larger enzymatic machinery for molecular functions such as gene transcription. Overall, these findings were consistent with the findings in A549 lung epithelial cells and BJ-fibroblasts indicating that packing domains may further generalize as the intermediate functional hierarchy of chromatin within human nuclei.

### RAD21 degradation depletes topological and loop domains in ensemble Micro-C

Owing to their physical size and the nucleotide composition within packing domains, we wanted to understand if (a) these structures reflected topological associated domains or loop domains and (b) if impairment of cohesion-mediated loop extrusion weakened the boundaries or sizes of packing domains proportional to the decrease observed on ensemble measures, such as Micro-C. To investigate the role of cohesin-mediated loop formation on packing domain structure, we utilized the AID2 system in HCT-116 cells to rapidly degrade endogenous RAD21 and analyzed publicly available high resolution Micro-C libraries available through the ENCODE Consortium as previously described (*12, 30–34*). We chose this cell line due to its superiority in preventing leakage-associated loss of the RAD21, the increased rapidity of degradation, and it’s decreased cytotoxicity(*30*). Given these features, analysis of its effect on genome structure could provide new insights on the role of the crucial protein, RAD21 in cell function. To verify depletion of RAD21 following 6 hours of 5-Ph-IAA treatment, RAD21 protein levels were quantified independent of the degron construct using confocal microscopy (**Fig. 3a, Fig. S2)** demonstrating complete depletion of RAD21 in >90% of the cell population by both immunofluorescence and mClover levels (**Fig. 3a**). Utilizing publicly available data through the ENCODE Consortium(*32–34*), we then analyzed the effect of RAD21 depletion on TADs, loop domains, compartments, and genome connectivity in Micro-C (accession numbers available in **Table S1**). As expected, RAD21 depletion results in the marked decrease of TADs and loops, with a visually apparent transformation in contacts genome wide (**Fig. 3b-h**)(*13, 15, 35, 36*). Aligning with previous results, we observed a robust decrease in compartment strength (**Fig. 3c, Fig. S3**) alongside increased compartmentalization and some compartment switching from A to B, which was previously seen in NIPBL mutant mice(*15, 37*). Consistent with prior reports demonstrating complete loss of TADs and loops, we observed that RAD21 depletion within RAD21-mAID-Clover CMV-osTIR1(F74G) cells was robust. In particular, we observed that ∼97% of all TADs were lost (**Fig. 3d,h, Fig. S3**). Likewise, we observed that ∼95% of all loops observed in the control condition were no longer present (**Fig. 3e-h, Fig. S3**).

**Fig. 3.**
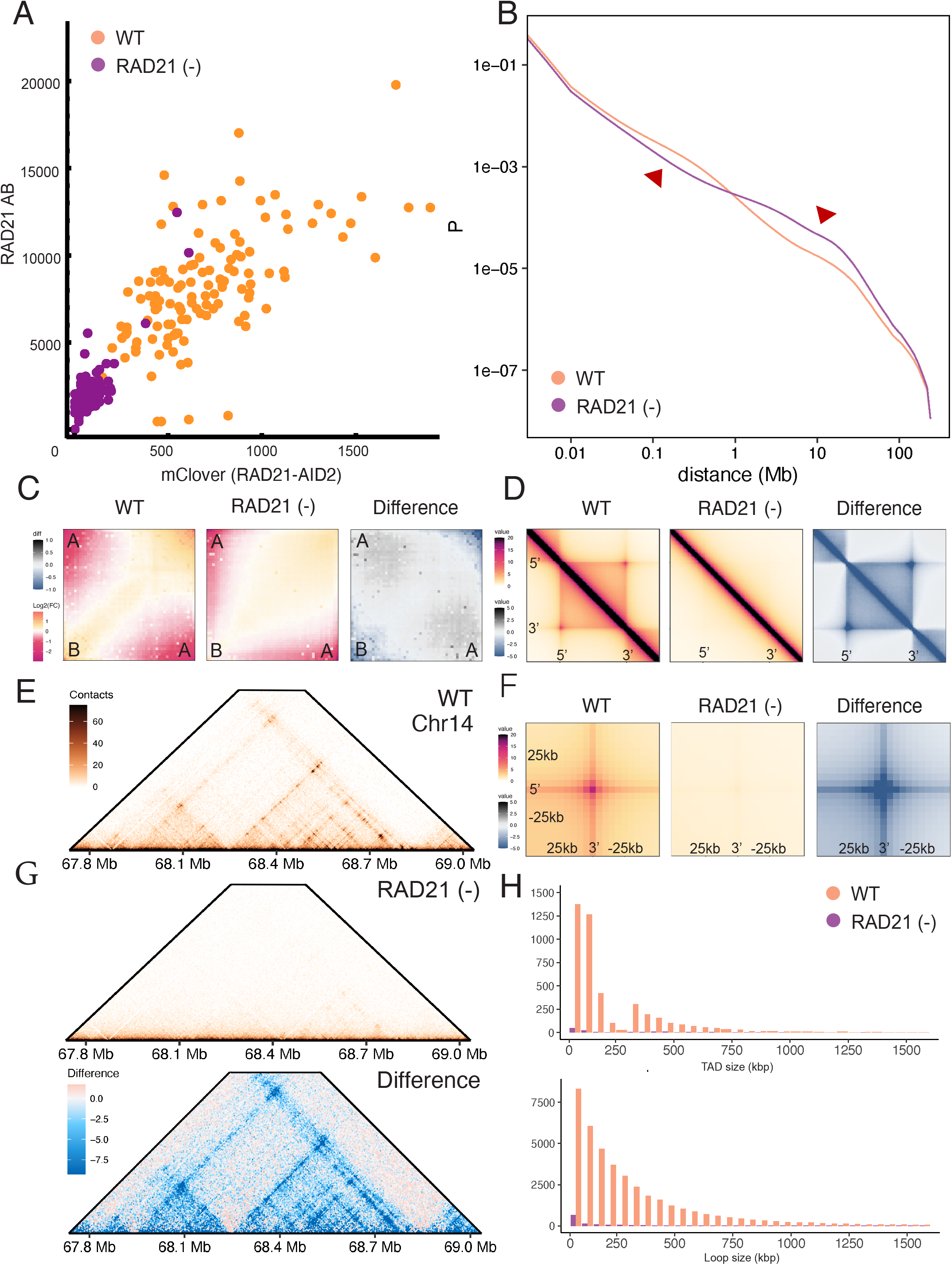
RAD21 depletion results in expected loss of TADs, loops, and decreased insulation. **(A)** Verification of RAD21 depletion by both quantification of mClover and RAD21-immunofluorescent staining showing over 90% of the population is without detectable RAD21. Axes representing corrected-total-fluorescence in arbitrary units. **(B)** Contact scaling is observed to decrease at short ranges and increase at long-ranges upon RAD21 depletion. **(C)** Compartment insulation demonstrates compartment score weakening with RAD21 depletion. (**D)** TAD insulation plots demonstrating TAD insulation strength decreases with RAD21-depletion. **(E)** Representative loop-domain anchor point on chromosome 14 showing that while the loop domain in control cells. **(F)** Observed weakening of loop-anchors on insulation plot. **(G)** Comparison of loop-domain observed in **(E)** upon RAD21 depletion demonstrating an decrease in local contacts. **(H)** Quantification of TADs and loop domains observed in control vs RAD21(-) cells showing predominantly a loss of >95% of TADs and ∼95% of loop domains at 6 hours of treatment with 1 μM 5-Ph-IAA.

### Packing domains largely persist despite RAD21 depletion

Having verified that (1) RAD21 was depleted throughout the population at short-time scales and (2) ensemble Micro-C demonstrated that TADs and loops were lost upon RAD21 depletion, we compared these findings to the structure of chromatin in ChromSTEM tomography of RAD21 cells treated with 1μM 5-Ph-IAA for 4 hours. In contrast to the findings on Micro-C, the ChromSTEM tomogram generated upon RAD21 depletion was visually very similar to those in DMSO-treated control cells, demonstrating a continuous distribution of heterogeneous structures and porosity throughout the examined field of view (**Fig. 4a**). Furthermore, the mass-scaling relationship observed in DMSO controls with the observation of three distinct chromatin regimes was not lost upon RAD21 depletion (**Fig. 4b**) and was comparable to that observed in the DMSO controls. With respect to both the disordered chromatin polymer (<25nm) and packing domains, RAD21 loss did not have a proportional effect on 3D genome organization to that observed on Micro-C. At the level of chromatin as a disordered polymer, we observed the existence of looping chromatin fibers (**Fig. 4c, Movie S3**) indicating that long range interactions persisted. Performing the analysis to identify packing domains as described above upon RAD21 depletion, we did observe an overall decrease in the number of packing domains (from 78 in DMSO treated cells to 62 in RAD21 depleted cells, ∼20% decrease) but the remaining domains were visually comparable to those observed in controls (**Fig. 4d, Movie S4**). Quantitatively, RAD21 loss only had a modest effect on the remaining packing domains, with small changes in CVC (0.40 → 0.43, p-val 0.20, **Fig. 4e**), *D* (2.61 → 2.60, p-val 0.45, **Fig. 4f**), and radius (84nm → 89nm, p-val 0.39, **Fig. 4g**). To test if these findings extended into live cells and were not due to transformation from fixation, we performed live-cell partial-wave spectroscopic (PWS) microscopy which can measure changes in *D* for chromatin between 20-200nm for control cells in comparison RAD21 depleted cells(*29, 38*). Comparable to the findings on ChromSTEM microscopy, we observed a minimal change in *D* in live cells upon 4 hours of RAD21 depletion (2.61 → 2.60, p-val = 0.104, **Fig. S4**) indicating that chromatin organization is predominantly preserved despite loss of RAD21 both on electron microscopy and in live-cells.

**Figure 4.**
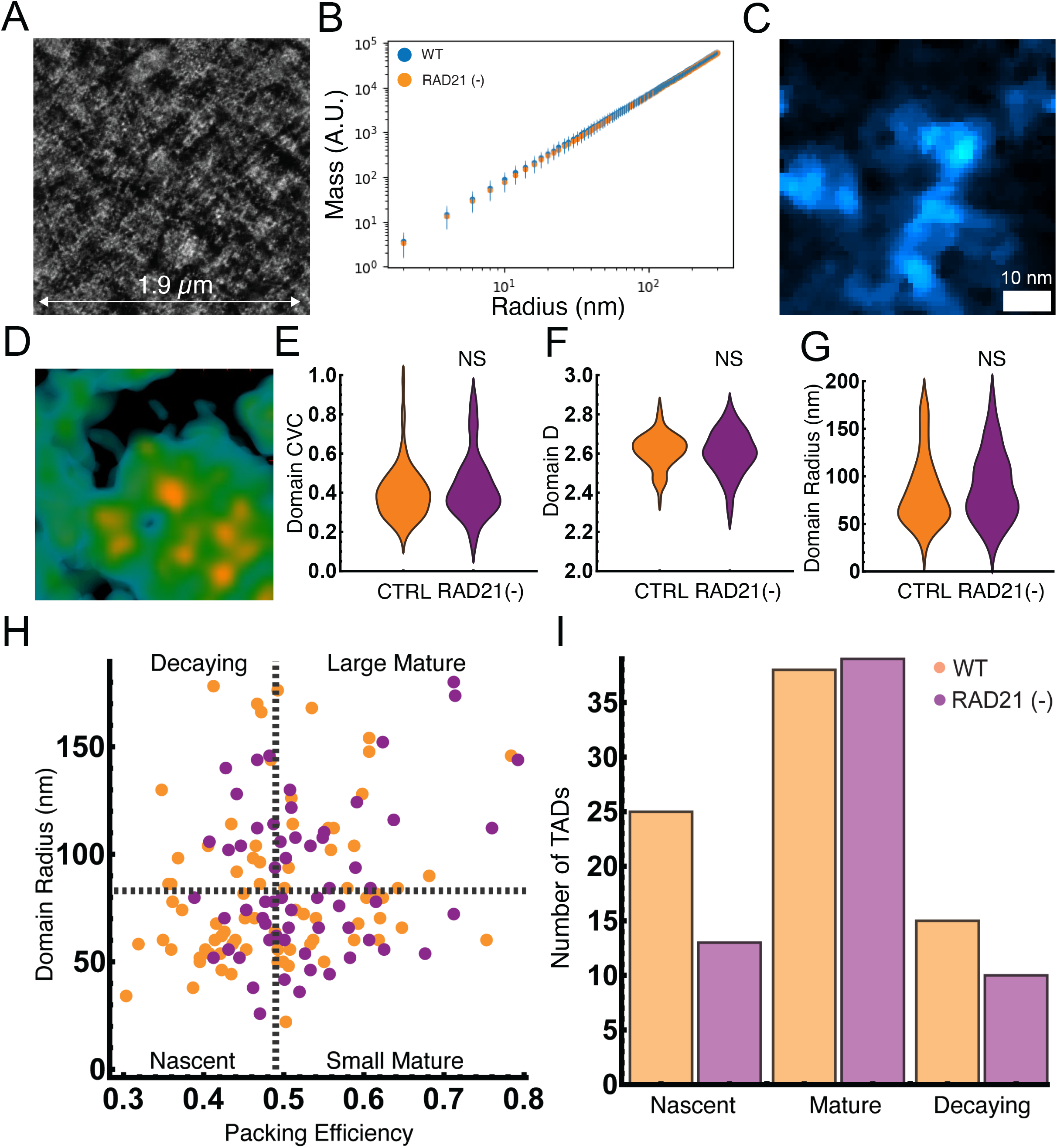
Packing domains size and organization are minimally transformed upon RAD21 depletion. **(A)** ChromSTEM tomogram from RAD21 5-Ph-IAA treated cells after 6 hours of depletion demonstrating remarkable similarity to chromatin DMSO control cells above. **(B)** Analysis of chromatin density distribution demonstrates minimal changes in chromatin organization across all three regimes. **(C)** Visualized chromatin loop in RAD21 depleted tomogram demonstrating continued presence of long-range interactions despite loss of cohesin-mediated loop extrusion. **(D)** Representative packing domain (200nm x 200nm) from RAD21 depleted cells showing similar features to those within control cells with a high-density center and continuous distribution of mass toward the periphery until the emergence of low density porous regions. **(E-G)** Overall, there was a total decrease in the number of observed packing domains from 78 to 62 upon RAD21 depletion, however, the remaining domains had similar CVC **(E)**, scaling of packing **(F)**, and size **(G)** in both conditions. P-values represent two-tailed unpaired t-test from the 78 and 62 packing domains. Bonferroni corrections were assumed for multiple comparisons (total 3) for an adjusted p-value of 0.0166. **(H)** Analysis of packing domains by size and packing efficiency to analyze domain properties. Nascent domains (low efficiency, small size), Mature domains (high packing efficiency), and Decaying domains (low efficiency, large size) are differentially impacted upon 1 μM ph-IAA for 6 hours. Black dashes represent mean packing efficiency and mean radius in control cells. **(I)** The primary decrease in packing domains occurred in nascent domains (low density, small domains; n = 25 to 13, ∼48% decrease) with a negligible change in the number of mature domains (high efficiency, n = 38 to 39, ∼2% increase). Decaying domains were also decreased (low efficiency, large size; n= 15 to 10, 33% decrease).

Although ChromSTEM imaging is restricted to fixed cells, in principle the observed chromatin structures capture a temporal cross-section of evolving structures. As such, one could consider that the size and density distribution of packing domains could relate to their current progress through a packing domain life-cycle. In such an analysis, denser structures represent mature domains whereas small, less dense structures represent nascent domains and finally large, low-density structures could represent decaying domains. As cohesin mediated looping could either be involved in barrier maintenance to prevent domain overlap or involved in domain formation(*20*– *22*), we analyzed the effect of RAD21 depletion on these subpopulations. We partitioned the features of packing domain across their size in comparison to their packing efficiency(*28*), which measures the per-unit area density of chromatin in the ChromSTEM tomogram (**Fig. 4h**). In this analysis, nascent structures are small and poorly packaged (below average size and below average packing-efficiency in control tomogram), mature structures can be large or small but are efficiently packed (above average efficiency), and decaying structures are large, poorly packed domains (above average size, below average efficiency). Notably, loss of RAD21 results in the largest change in the proportion of nascent domains (n=13 vs n=25 in controls, ∼48% decrease in number) whereas there was a minimal effect in the number of mature domains (n=39 vs n=38 in controls, ∼2% increase, **Fig. 4h,i**). In the context that RAD21 is depleted within 1 hour in this cell-line model(*30*) (**Fig. S5**), this indicates that mature domains do not depend on cohesin function for continued stability. Further, that half of nascent domains were still observed at 4 hours indicates the presence of a secondary mechanism that induces domain formation. It is also worth noting that ChromSTEM imaging captures the statistics of single cell organizational states, whereas the ensemble measurement of Hi-C is an aggregate statistic of many organizational states. Consequently, this suggested that mature packing domains were largely maintained independent of cohesin mediated loop formation but that the principal effect of RAD21 extrusion was on the production of a subset of nascent domains, possibly consistent with findings in prior reports using oligo-based methods(*20, 22*).

The retention of the majority (∼80%) of supra-nucleosome packing domains contrasted sharply with the loss of ∼97% of TADs with RAD21 depletion (**Fig. 3d-h, Fig 4h,i**) and 95% of loop domains (**Fig. 3d-h, Fig 4h,i**)) suggested it was possible that these features do not map directly to a space-filling physical entity but instead serve to coordinate the retention of distant loci in close proximity with additional mechanisms determining their local physical configuration. To test this hypothesis, we first compared the DNA content contained within TADs, loop domains, and packing domains, and found a similar distribution of genomic content in both structures (**Fig. 5a**). Specifically, the genomic content within packing domains sizes ranged between 6kbp to 1.3mbp in control cells (median size of 89 kbp) in comparison to TADs with a range of 60kbp to 2.5mbp (median size 130kbp), and loop domains ranging from 9kbp to 9 mbp (median size of 170kbp). We hypothesized that if TADs or loop domains represented high frequency packing domains, that their loss would result in changes in accessibility as these structures would no longer have the space-filling features observed in packing domains of dense cores with accessible surfaces (**Fig. 4c**). Likewise, if global alterations in connectivity resulted in genome-wide alterations in accessibility, we would observe throughout the genome an increase in accessible loci consistent with the loss of numerous space-filling structures. To test this hypothesis, we utilized publicly available ATAC-Seq data from the ENCODE Consortium(*32–34*) on RAD21 depleted AID2 cells (accession numbers in **Table S2**), finding no change in the frequency of accessible loci per chromosome with or without RAD21 depletion at 6 hours (**Fig. 5b**), suggesting that loss of loop domains does not result in global alterations in chromatin accessibility. Next, we tested whether local accessibility within TAD loci or loop domains would increase, which would be consistent with a shift in local access upon the loss of these structures if they represented physical structures. Contrary to the hypothesis that TADs and loop domains are space-filling features of the genome, we again found no change in the local accessibility in either TADs (**Fig. 5c**) or loop domains (**Fig. 5d**) upon RAD21 depletion. Overall, these findings were consistent with our results in ChromSTEM as most domains were not transformed with RAD21 depletion. Paired with prior work showing that heterochromatin domains are stable after RAD21 loss(*25*), it is more likely that TADs and loop domains are representative measures of population genomic connectivity and may not directly map to space-filling structures observable on ChromSTEM.

**Fig. 5.**
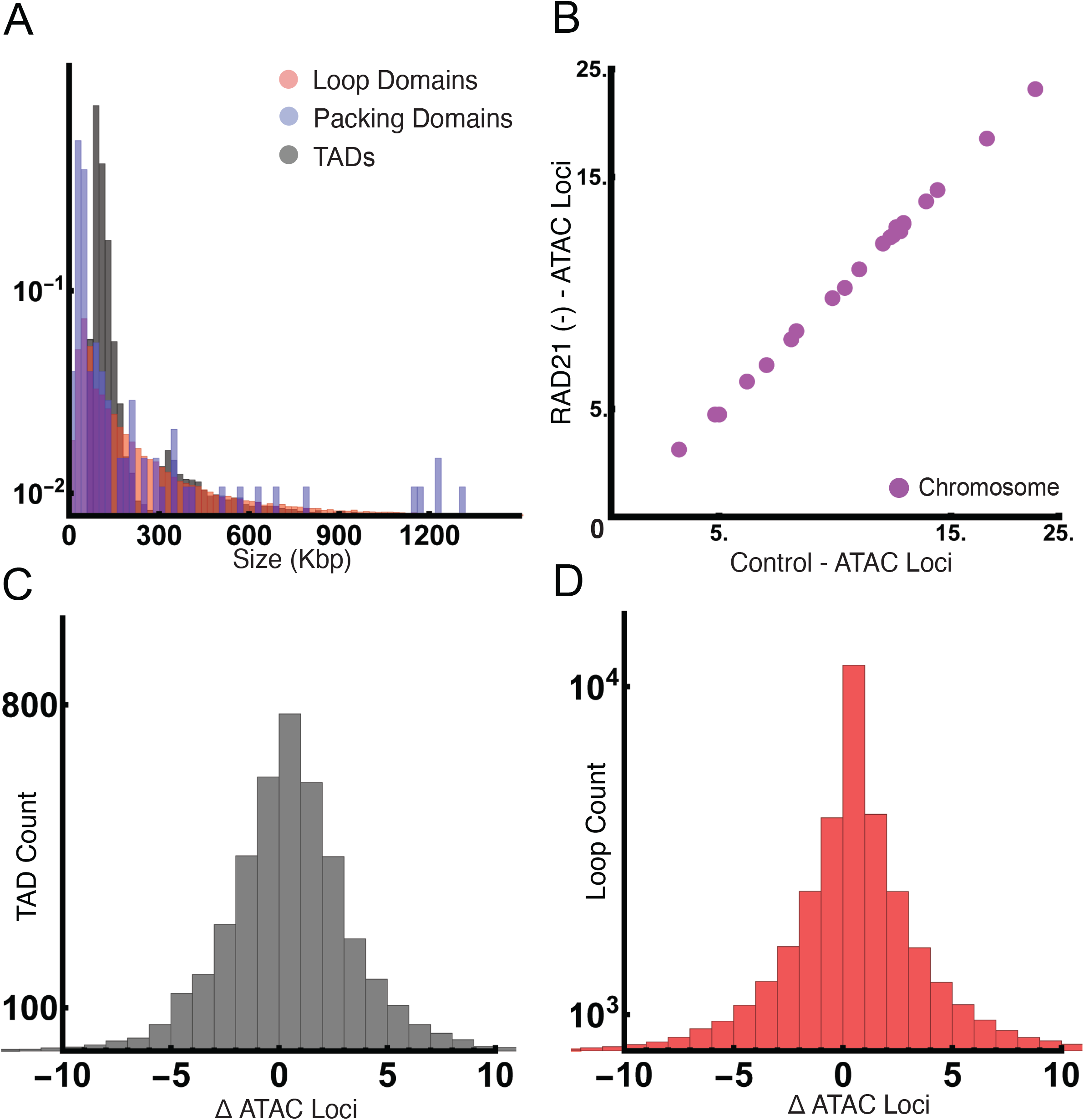
Analysis of TAD and loop domains indicates no significant change in global or local accessibility with cohesin depletion. **(A)** Comparison of packing domain DNA content (median 89.32 kbp, minimum of 6.27kbp and max of 1.306mbp) in comparison to loop domains (median 170kbp, minimum of 9kbp and max of 9.06Mbp) and TADs (median 130kbp, minimum 60kbo and max of 2.55Mbp) showing similar ranges in size in control cells. **(B)** Analysis of ATAC-Seq accessibility per chromosome in control cells vs. RAD21(-) cells showing no significant change in global accessibility. Axes are reported in number of loci times 10^3^. **(C)** Analysis of the change in number of ATAC-Seq loci within TAD coordinates before and after RAD21 depletion showing no change in accessibility in TAD regions upon their loss (median = 0, interquartile range of –2 to +2). **(D)** Analysis of the change in number of ATAC-Seq loci within loop domain coordinates before and after RAD21 depletion showing no change in accessibility in loop regions upon their loss (median = 0, interquartile range of –2 to +2).

## Discussion

Chromatin electron microscopy maps DNA density proportionally to the measured contrast on imaging, resolving both the space-filling properties of the genome and its polymeric structure in 3D. Critically, ChromSTEM identifies a transition topology of chromatin from a disordered polymer (<25nm) into higher-order structures that include packing domains and chromatin territories(*27–29*). By resolving the structures directly, it is evident that genome organization includes biphasic, porous packing domains (**Fig. 1c**) and chromatin loops (**Fig. 1d**) *in vitro*. In the context of prior work, the previously observed structures of chromatin are now evident in three distinct cell line models: A549 cells (lung adenocarcinoma epithelial cells), HCT-116 cells (microsatellite unstable colonic epithelial cells), and BJ cells (immortalized fibroblasts). Although fixation alone can alter chromatin structure, prior work demonstrated that the ChromSTEM preparation protocol still maintains structural concordance with cells imaged via live-cell nanoscopy through the sample preparation stages(*29, 38–40*). When combined with the similar findings observed using live cell PWS microscopy (**Fig. S4**) that *D* is unchanged upon RAD21 depletion, the resultant ChromSTEM findings appear concordant with the organization of chromatin in live-cells.

The major focus of this work is to understand if cohesin-mediated loop formation is the primary mechanism behind generating and maintaining chromatin packing domains. We investigated the effect of cohesin depletion on packing domains given that their size is approximately that previously reported to represent TADs and loop domains (**Fig. 5a**). We observed that RAD21 depletion results in the profound decrease in TADs and loops (**Fig. 3**) but does not result in a proportional change in packing domain frequency, with only a decrease of ∼20% of packing domains. As most domains persist after depletion while retaining their baseline structure (similar size, genomic content, and polymeric properties), this indicates that most packing domains are sustained independent of cohesion-mediated loop extrusion (**Fig. 4**). Although this study has limitations in that it measures packing domains in sub-portion of one nucleus in each group, other studies have similarly showed the presence of TAD-like structures after cohesin degradation(*17, 20–22, 25*). While it is possible that TAD-like structures could be an observed subset of chromatin packing domains, the observed discordance between the number of lost TADs and the retention of most domains suggests otherwise. In support of these findings, we observe minimal changes in global and local accessibility upon RAD21 depletion on ATAC-Seq, indicating TADs and loops are measures of connectivity but may not impart direct barrier function (**Fig. 5b-d**). Ultimately, future work investigating if previously observed TAD-like structures translate into packing domains will require further technological development including the ability to identify specific genomic loci on electron microscopy while maintaining proportional mapping of stain density to DNA content without degradation of nanoscale structure(*26*).

That packing domains are largely unaffected by the disruption of RAD21 (**Fig. 4**) offers several interesting insights into genome organization. For example, prior work has shown that gene transcription itself could produce long-range promoter-enhancer interactions forming ‘loops’ that would be one additional mechanism that could result in domain formation(*25, 41*). Likewise, packing domains could arise from the enzymatic activity of heterochromatin remodelers or other forces such as ionic charge shielding that could increase monomer-monomer interactions of a polymer to form long range interactions(*25, 42*). Alternatively, it is possible that packing domains do not emerge from a single mechanism, but instead arise from the confluence of factors that govern both long range interactions (loop extrusion, promoter-enhancer looping) with mechanisms that modify monomer-monomer interactions (histone methylation or acetylation, ionic shielding) like what was demonstrated in synthetic chromatin assembly(*43, 44*). The investigation of such phenomenon in future works could expand our understanding of the molecular and physiochemical regulatory mechanisms of chromatin organization.

## Supporting information

Control domain structure

control loop structure

rad21 depleted domain structure

rad21 depleted loop structure

## Acknowledgments

Chat-GPT 4.0 was utilized to assist in the review of the logic-structure in the manuscript and with coding support for data visualization. No AI tools or external reviewers were utilized for conceptualization, writing, synthesis, or direct analysis of results. Philanthropic support was generously received from Rob and Kristin Goldman, the Christina Carinato Charitable Foundation, Mark E. Holliday and Mrs. Ingeborg Schneider, and Mr. David Sachs. Computational analysis of Hi-C data was supported in part through the computational resources and staff contributions provided by the Genomics Compute Cluster, which is jointly supported by the Feinberg School of Medicine, the Center for Genetic Medicine, and Feinberg’s Department of Biochemistry and Molecular Genetics, the Office of the Provost, the Office for Research, and Northwestern Information Technology. The Genomics Compute Cluster is part of Quest, Northwestern University’s high-performance computing facility, with the purpose to advance research in genomics. We appreciate the generous support from the ENCODE Consortium in the generation and dissemination of publicly available datasets. In particular, we thank the lab of Dr. Erez Lieberman-Aiden for the generation of the Micro-C dataset and the lab of Dr. Michael Snyder for generation of the ATAC-Seq dataset in RAD21 depleted and control cells with the AID2 system.

## Funding

National Science Foundation grant EFMA-1830961 (EPL, MC, IS, VB)

National Science Foundation grant EFMA-1830969 (EPL, VB)

National Institutes of Health grant R01CA228272 (WSL, TK, IS, VB)

National Institutes of Health grant U54 CA268084 (WSL, LMC, LMA, EPL, TK, KLM, MC, IS, VB)

National Institutes of Health grant U54 CA261694 (LMC, VB)

NIH Training Grant T32AI083216 (LMA)

NIH Training Grant T32GM132605 (TK, LMC, VB)

Hyundai Hope on Wheels Hope Scholar Grant (KLM)

CURE Childhood Cancer Early Investigator Grant (KLM)

Alex’s Lemonade Stand Foundation ‘A’ award grant (KLM)

Ann and Robert H. Lurie Children’s Hospital of Chicago under the Molecular and Translational Cancer Biology Neighborhood (KLM)

## Author contributions

Conceptualization: LMA, MC, MTK, IS, VB

Formal Analysis: WSL, LMC, LMA, EP, TK

Methodology: VD, MTK, IS, VB

Investigation: WSL, LMC, LMA, EP, TK, KLM

Visualization: WSL, LMC, LMA, EP, TK

Funding acquisition: VD, MTK, IS, VB

Project administration: LMA, VB

Supervision: LMA, VD, MTK, IS, VB

Writing – original draft: WSL, LMC, LMA, EP

Writing – review & editing: WSL, LMC, LMA, EP, TK, KLM, MC, VD, MTK, IS, VB

## Data and materials availability

All of the data, analysis software, and relevant materials have been uploaded to the following locations. Micro-C and ATAC-Seq data were obtained from the ENCODE Consortium with the relevant accession numbers posted in the supplementary information in Tables S1 and S2. Analysis software and relevant compiled data to regenerate the produced figures has been uploaded to GitHub at github.com/BackmanLab/RAD21. Raw and processed ChromSTEM-HAADF tomograms are available from the authors upon request.

