## Supplemental Materials and figures for "Chromatin packing domains persist after RAD21 depletion in 3D"

**The PDF file includes:**

#### Supplementary Materials:

Materials and Methods

Figs. S1 to S5

Table S1 to S3

### Materials and Methods

#### Cell Culture

HCT116 cells (ATCC, #CCL-247) and HCT116 RAD21-mAID-Clover CMV-OsTIR1(F74G) cells were grown in McCoy's 5A Modified Medium (#16600-082, Thermo Fisher Scientific, Waltham, MA). All cell culture media was supplemented with 10% FBS (#16000-044, Thermo Fisher Scientific, Waltham, MA) and penicillin-streptomycin (100 μg/ml; #15140-122, Thermo Fisher Scientific, Waltham, MA). Cells were maintained under recommended conditions at 37°C and 5% CO2. Cells were allowed at least 24 hours to re-adhere and recover from trypsin-induced detachment. All imaging was performed when the surface confluence of the dish was between 40–70%. All cells in this study were maintained between passage 5 and 20. All cells have been tested for mycoplasma contamination (ATCC, #30-1012K) before starting experiments, and they have given negative results. To create HCT116 RAD21-mAID-Clover CMV-OsTIR1(F74G) cells, HCT116 cells (ATCC, #CCL-247) were modified with the AID system as previously described(*29*, *30*). Briefly, when the *Oryza sativa* TIR1 (OsTIR1 (F74G) mutant is expressed in non-plant cells, it forms a Skp1–Cul1–F-box (SCF) E3 ligase complex with endogenous components. In the presence of 5-phenyl-indole-3-acetic acid (5-Ph-IAA), the protein of interest (RAD21) that is fused to a 7 kDa degron (mini-AID) is rapidly degraded rough the ubiquitin–proteasome pathway(*29*, *30*).

#### 5-Ph-IAA Treatment

HCT116 RAD21-mAID-Clover CMV-OsTIR1(F74G) cells were plated at 50,000 cells per well of a 6-well plate (Cellvis, P12-1.5H-N). To induce degradation of RAD21, 5-Ph-IAA (#HY-134653, MedChemExpress), 5-(3,4-dimethylphenyl)-indole-3-acetic acid, 5-(3-methylphenyl)-indole-3-acetic acid and 5-(3-chlorophenyl)-indole-3-acetic acid was dissolved in DMSO to make a 500 mM stock solution, and further diluted with DMSO to make a working stock solution of 1 mM immediately before the experiment. A final concentration of 1 μM of 5-Ph-IAA was added to HCT116 RAD21-mAID-Clover CMV-OsTIR1(F74G) cells for 6 hours(*29*, *30*).

#### ChromSTEM-HAADF

*Electron Microscopy Sample Preparation*

Samples were prepared as previously described(*26–28*). Reagents used are summarized in Table S3. The cells were fixed with 2% paraformaldehyde, 2.5% glutaraldehyde (EM-Grade), 2mM $CaCl_2$ in 0.1M sodium cacodylate buffer for 30 minutes in room temperature and 30 minutes in fridge. The cells were then kept in cold temperature, if possible, for further treatments. After fixation, the cells were washed 5 x 2 minutes with 0.1M sodium cacodylate buffer and blocked with 10mM glycine, 10mM potassium cyanide, 0.1M sodium cacodylate buffer for 15 minutes.

The cells were stained with 10μM DRAQ5, 0.1% saponin, 0.1M sodium cacodylate buffer for 10 minutes, followed by 3 x 5 minutes washing with the blocking buffer. After that, the cells were photo-oxidized in 2.5mM 3,3'-Diaminobenzidine (EM-Grade) under a 100X oil objective, 15W Xenon lamp and Cy5 filter for 5 minutes. The cells were washed 5 x 2 minutes with 0.1M sodium cacodylate buffer and stained with 2% osmium tetroxide, 1.5% potassium ferrocyanide, 2mM $CaCl_2$ in 0.15M sodium cacodylate buffer for 30 minutes. The cells were washed 5 x 2 minutes with Millipore water afterwards.

The cells were then dehydrated gradually with 30%, 50%, 70%, 85%, 95%, 2 times 100% ethanol. After that, the cells were incubated under room temperature with 100% ethanol, and infiltrate and embed with Durcupan resin with standard procedures. After 48 hours of resin incubation at 60$^o$c, the resin blocks were collected for ultramicrotomy.

For ultramicrotomy, an ultramicrotome (UC7, Leica) and a 35$^o$ Diatome knife were used to section 120nm thick resin samples. The sections were collected on a copper slot grid (2 x 0.5mm) with formvar/carbon film. Gold nanoparticles with 10nm diameter were deposited on both surfaces of the grid afterwards as fiducial markers.

*Image Collection and Tomography Reconstruction*

Images were collected with Hitachi HD2300 STEM microscope at 200kV with HAADF imaging mode, at a magnification of 50kX. Two sets of tilt series images were collected by rotating samples from -60$^o$ to +60$^o$ at a 2$^o$ step, with two roughly perpendicular rotation axes.

For post-processing, IMOD was used to align the images. The gold nanoparticles of the collected images were removed with IMOD for another set of data without the influence of extreme values from gold nanoparticles. For each tilt series, Tomopy was used to reconstruct the volume with penalized maximum likelihood algorithm with weighted linear and quadratic penalties. The two independent reconstructed volumes were then combined in IMOD, with gold nanoparticles as the matching model and repeated on the data without nanoparticles.

After reconstruction, the top and the bottom 0.1% pixel values were capped to remove extreme values. The pixel values are then scaled between 0 to 1 for analysis.

*Chromatin Domain Identification and Analysis*

Chromatin domains were identified and analyzed following the approach previously mentioned(*27*, *28*).

For identification of domains, a Gaussian filter with radius = 5 pixels was used followed by CLAHE contrast enhancement on the 2D projection of the 3D tomogram using ImageJ. Chromatin domain centers were identified as local maxima with prominence = 1.5 x standard deviation of pixel values.

For domain properties, an 11 x 11 pixels window is applied to each domain and sampled for each domain. Packing scaling analysis is done by measuring the total intensity of the chromatin that radially expands from the center pixel picked and weighted by the intensity of the center pixel. The linear region of the packing scaling behavior is identified by MATLAB 'ischange' function. The domain size is measured as the point at which the packing scaling behavior deviates from the linear fits with 5% difference, or when local packing scaling exponent D reaches 3. CVC of a domain is measured with the average value within the domain on a binarized image with Otsu-binarization algorithm after CLAHE contrast enhancement in ImageJ. The packing efficiency, $A$, is calculated as previously described from the relationship described previously(*27*, *28*). Briefly, $I = A*(R_f/R_{min})^D$, where I is the average chromatin intensity of a domain, $R_f$ is the size of the domain, $R_{min}$ = 10nm is the smallest unit of random chromatin polymer chain and $D$ is the packing scaling exponent of the domain packing behavior.

Loop images and videos were generated from the produced tomograms and then underwent background subtraction with an applied local filter for visualization.

### Micrococcal Nuclease Chromatin Conformation Capture

#### Micro-C data processing and analysis:

Processed Micro-C files were obtained from publicly available data through the ENCODE Consortium (**Table S1**)(*31–33*). TAD and loop domains were obtained from the reported bedpe files (**Table S1**). Compartment eigenvector analysis and Pearson correlation analysis were generated using Juicer's Eigenvector and Pearsons scripts, respectively, or using built in functions in GENOVA. Prior to analysis, Hi-C contacts were dumped using Juicer's Straw (https://github.com/aidenlab/straw). Contacts were converted to coolers and normalized using Cooler (https://github.com/open2c/cooler) at 5kb and 100kb resolution. Aggregate TAD Analysis and Aggregate Peak Analysis were generated on these contact maps using GENOVA (https://github.com/robinweide/GENOVA). Visualization of contact loci was also done using GENOVA. All other analyses were custom-generated in R. All code used in this publication is available on Github at: https://github.com/BackmanLab.

### Assay for transposase-accessible chromatin with high throughput sequencing (ATAC-Seq)

Processed ATAC-Seq bed files were obtained from publicly available data through the ENCODE Consortium (**Table S2**)(*31–33*).Analysis of global accessibility was performed by analyzing the per-chromosome number of accessible loci previously identified with a q-value threshold greater than 1. The location of each peak was approximated by the average position reported between the 3' and 5' locations. Analysis of accessibility within TADs and loop domains was performed by utilizing the indices reported above (**Table S1**). For loop domains, the mean 3' and 5' end for each anchor loci were calculated and used for subsequent analysis of accessibility. Within each individual TAD or loop location, the relative accessibility was calculated by measuring the change in the number of peaks within each TAD or loop domain after RAD21 depletion.

### Live-cell Partial Wave Spectroscopic (PWS) Microscopy

For live-cell measurements, cells were imaged and maintained under physiological conditions (5% $CO_2$ and 37°C) using a stage-top incubator (In Vivo Scientific, Salem, SC; Stage Top Systems). The PWS optical instrument was built on a commercial inverted microscope (Leica, Buffalo Grove, IL, DMIRB) supplemented with a Hamamatsu Image-EM CCD camera C9100-13 coupled to an LCTF (CRi Woburn, MA) for hyperspectral imaging. Spectrally resolved images of live cells were collected between 500 and 700 nm with a 2-nm step size(*28, 37*). Broadband illumination was provided by an Xcite-120 light-emitting diode lamp (Excelitas, Waltham, MA). PWS is a high-throughput, label-free technique that measures the spectral standard deviation ($\Sigma$) of internal optical scattering originating from chromatin(*37*). Variations in the refractive index distribution $\Sigma$, are characterized by a mass density autocorrelation function (ACF) to calculate chromatin packing, scaling $D$(*28*). Changes in $D$ resulting from each condition are quantified by averaging cells, taken across 3 technical replicates. Population $D$ are calculated by first averaging $D$ values from PWS measurements within each cell nucleus and then averaging these measurements over the entire cell population for each treatment condition. Since $D$ is uncorrelated from cell to cell except for direct progeny, statistical comparison is performed across the whole population instead of on the technical replicates(*28*).

### Western Blot Analysis

Total cellular protein from HCT116 RAD21-mAID-Clover CMV-OsTIR1(F74G) cells was extracted using the Western-Ready Rapid Protein Extraction Buffer (BioLegend, #426305) following the manufacturer's protocol. Cell lysates were quantified with a standard Bradford assay using the Pierce Bradford Plus Protein Assay Reagent (Thermo Fisher Scientific, #23238) and Pierce Bovine Serum Albumin Standard Pre-Diluted Set (Thermo Fisher Scientific, #23208). Heat denatured protein samples were resolved on a Bolt Bis-Tris Plus Mini Gel 4–12% (Invitrogen, # NW04125BOX) and transferred to a PVDF membrane using the iBlot 2 Gel Transfer Device (Invitrogen, #IB21001) (20V for 7 minutes). Antibodies were bound to the membrane using the iBind Flex Western System (Invitrogen, # SLF2000) and associated reagents. Whole-cell lysates were blotted against the following primary antibodies: anti-RAD21 (Abcam, #ab217678, dilution 1:200) and anti-GAPDH (Sigma Aldrich, # G9545, dilution 1:1000). The following secondary antibody was used: Goat anti-Rabbit IgG (Heavy chain), Superclonal Recombinant Secondary Antibody, HRP (Invitrogen, #A27036, dilution 1:4000). To develop blots for protein detection, SuperSignal West Pico PLUS Chemiluminescent Substrate (Thermo Fisher Scientific, #34580) was used. To quantify the western blot bands, we used the iBright CL1500 Imaging System (Invitrogen, #A44240) and iBright Analysis Software to define bands as regions of interest. Samples were loaded in duplicate on the same gel. The blot was then sectioned in half after transfer to facilitate detection of GAPDH and RAD21.

Samples were loaded in duplicate on the same gel. The blot was then sectioned in half after transfer to facilitate detection of either RAD21 or GAPDH

### Statistical Analysis

Statistical analysis was performed using GraphPad Prism 10.1.1, Microsoft Excel, and Mathematica. Pairwise comparisons were calculated on datasets consisting of, at a minimum, biologically independent duplicate samples using two-tailed unpaired t test. The type of statistical test is specified in each case. A P value of $< 0.05$ was considered significant. Statistical significance levels are denoted as follows: n.s. = not significant; *$P<0.05$; **$P<0.01$; ***$P<0.001$; ****$P<0.0001$. Sample numbers (# of nuclei, n) or packing domains and the type of statistical test used are indicated in figure legends. Corrections for multiple comparisons on the same data were considered and Bonferroni correction applied as indicated where appropriate.

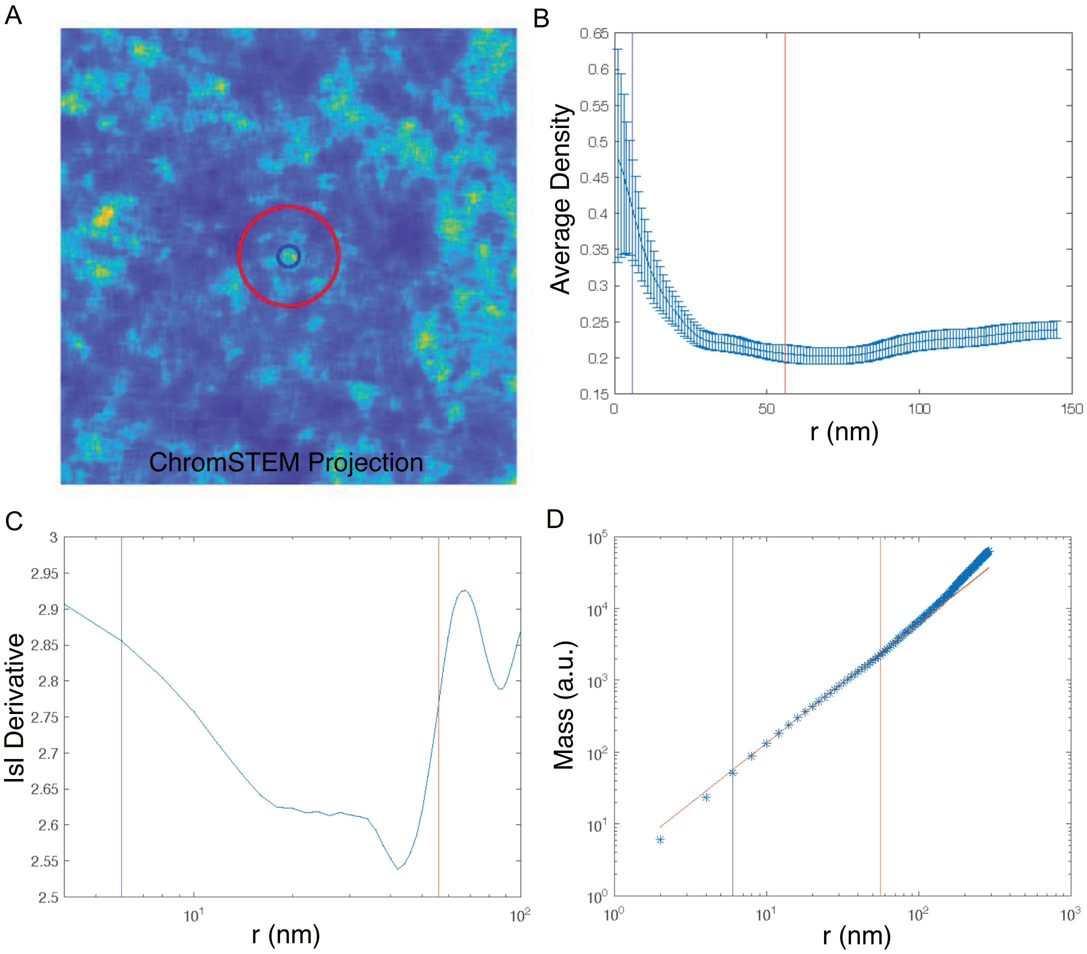


**Fig. S1. Identification of packing domain boundaries. A)** Representative packing domain center with surrounding calculated radius. **B)** Analysis of the radial distribution of mass from the center of the domain to identify the boundary. The boundary by this method is defined by the point where the mass is comparable to the average mass throughout the tomogram. **C)** Identification of domain boundary by calculating the point where the first derivative of the mass scaling equals 3. **D)** Analysis of domain radius by the radial density distribution function. The boundary by this method is defined by the radius where the mass deviates from power-law scaling (deviation from the log-log fit). The minimum radius from these three methods is then defined as the boundary of the packing domain.

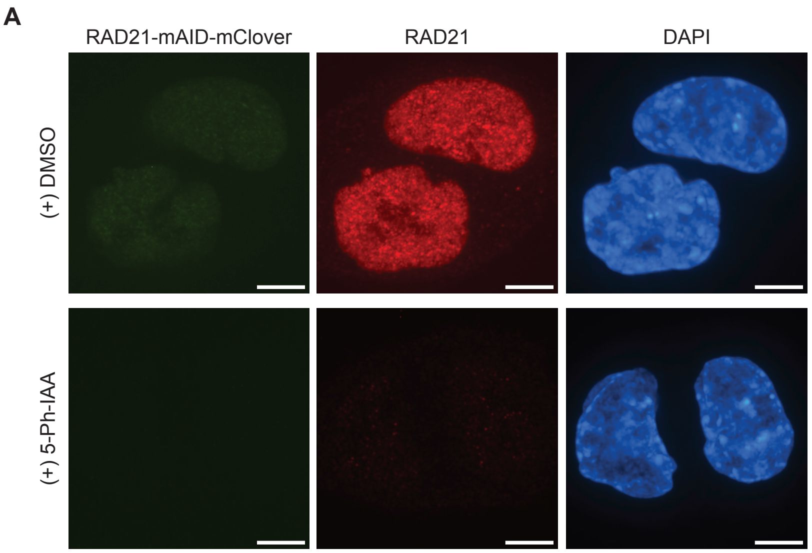


**Fig. S2. Representative images of DMSO controls and RAD21(-) at 6 hours.** Representative fluorescence image of control cells vs RAD21 depleted cells at 6 hours. First column represents mClover signal, second column represents immunofluorescence signal from RAD21 antibody staining, third column represents DAPI nuclear counter staining. Scale bar represents 5 microns.

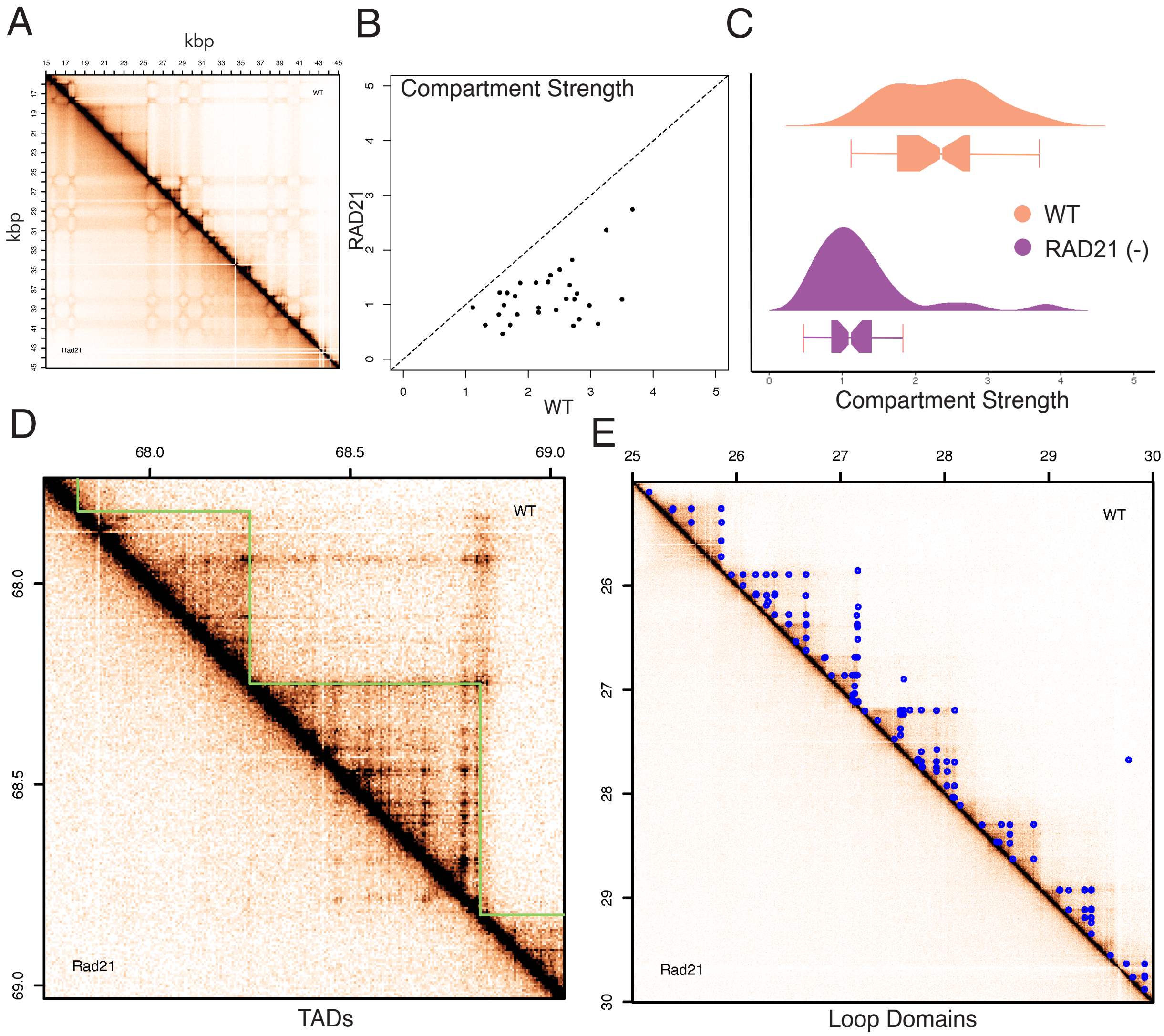


**Fig. S3. Extended analysis of DMSO control and RAD21(-) Micro-C features. A)** Representative change in compartments upon RAD21 depletion. **B)** Alterations in compartment strength per chromosome. **C)** Genome-wide transformation of compartment strength. D) Representative TAD loci before and after RAD21 depletion. **E)** Representative loop domains before and after RAD21 depletion

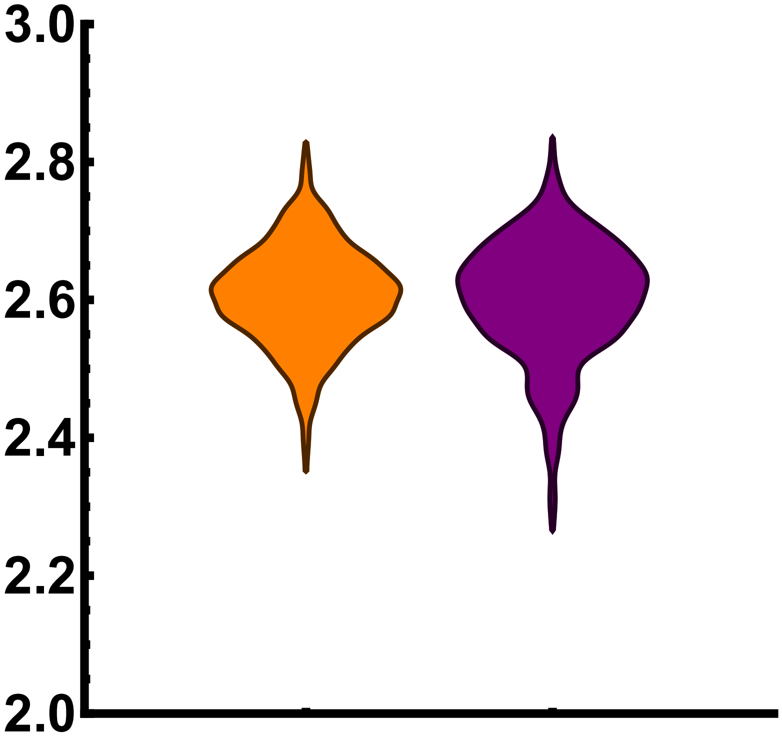


**Fig. S4 Live-cell Partial Wave Spectroscopic imaging of HCT116 cells, DMSO control, and RAD21(-) cells.** Comparison of chromatin organization in DMSO treated control cells (orange, n=916 cells from 3 replicates; mean *D* of 2.61 with a standard deviation of 0.072) in comparison to 5-Ph-IAA treated cells at 4 hours (purple, n = 922 from 3 replicates, mean *D* of 2.60 with standard deviation of 0.086). Two-tailed, unpaired T-test with p-value of 0.104.

### Uncropped Western Blots

**1st Replicate** **2nd Replicate**

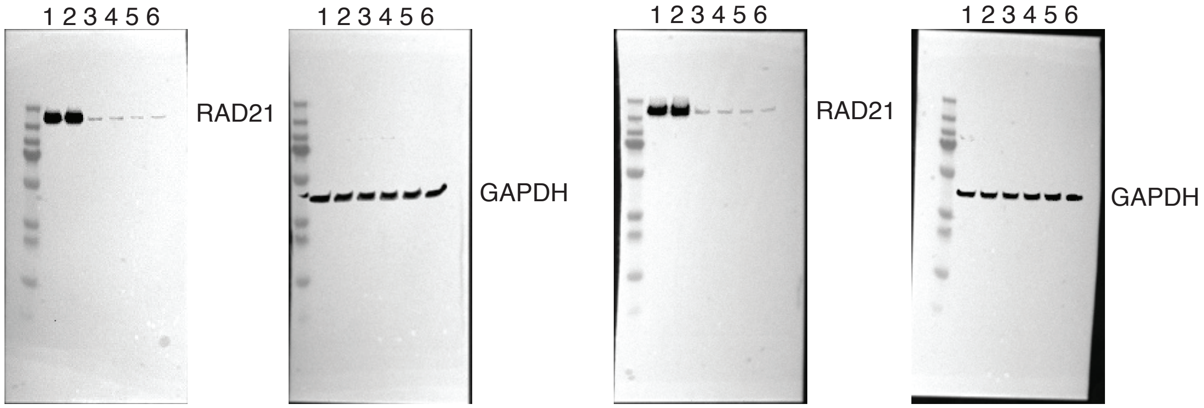


**NOTES:**

- 6 µg loaded per well of HCT116 RAD21-mAID-Clover CMV-OsTIR1 (F74G) lysate
- Order of Lanes:
  1. Untreated
  2. 6 hour DMSO
  3. 1 hour 1µM 5-Ph-IAA
  4. 2 hour 1µM 5-Ph-IAA
  5. 4 hour 1µM 5-Ph-IAA
  6. 6-hour 1µM 5-Ph-IAA
- RAD21 bands at ~135 kDa
- GAPDH bands at ~37 kDa
- **Ladder:** PageRuler Plus Prestained Protein Ladder (Thermo Fisher Scientific, Cat: 26619)

**Fig. S5 Western Blot analysis of RAD21 depletion over time after 5-Ph-IAA treatment.** Rapid depletion of RAD21 is achieved within 1 hour of exposure to 1 micromolar of 5-Ph-IAA in HCT RAD21-mAID-Clover CMV-OsTIR1(F74G) cells.

### Table S1. Micro-C ENCODE Consortium Files Utilized

| Experiment | File | Method | Source | Type |
|---|---|---|---|---|
| **ENCSR958BEA** | ENCFF794OIH | Micro-C | Genetically modified HCT-116 cell lines with CRISPR inserting O. sativa LOC4335696. Untreated. | Bedpe file; loop domains |
| **ENCSR958BEA** | ENCFF334LVK | Micro-C | Genetically modified HCT-116 cell lines with CRISPR inserting O. sativa LOC4335696. Untreated. | Bedpe file; contact domains |
| **ENCSR958BEA** | ENCFF528XGK | Micro-C | Genetically modified HCT-116 cell lines with CRISPR inserting O. sativa LOC4335696. Untreated. | .hic file; contact matrix |
| **ENCSR087JOM** | ENCFF630HHH | Micro-C | Genetically modified HCT-116 cell lines with CRISPR inserting O. sativa LOC4335696 treated with 5-Ph-IAA for 6 hours for RAD21 depletion | Bedpe file; loop domains |
| **ENCSR087JOM** | ENCFF307DPS | Micro-C | Genetically modified HCT-116 cell lines with CRISPR inserting O. sativa LOC4335696 treated with 5-Ph-IAA for 6 hours for RAD21 depletion | Bedpe file; contact domains |
| **ENCSR087JOM** | ENCFF317OIA | Micro-C | Genetically modified HCT-116 cell lines with CRISPR inserting O. sativa LOC4335696 treated with 5-Ph-IAA for 6 hours for RAD21 depletion | .hic file; contact matrix |

**Table S2. ATAC-Seq ENCODE Consortium Files Utilized**

| Experiment | Bed File | Method | Source |
|---|---|---|---|
| **ENCSR135OML** | ENCFF460JUY | ATAC-Seq | Genetically modified HCT-116 cell lines with CRISPR inserting O. sativa LOC4335696 treated with 5-Ph-IAA for 6 hours for RAD21 depletion |
| **ENCSR389WJO** | ENCFF107INQ | ATAC-Seq | Genetically modified HCT-116 cell lines with CRISPR inserting O. sativa LOC4335696 |

**Table S3. Reagents used in ChromSTEM sample preparation**

| Reagent | Formula |
|---|---|
| Washing solution | Hank's balanced salt solution without calcium and magnesium |
| Fixation solution | 2.5% EM grade glutaraldehyde<br>2% paraformaldehyde<br>2 mM $CaCl_2$<br>0.1 M sodium cacodylate buffer, pH = 7.4 |
| Blocking solution | 10 mM glycine<br>10 mM potassium cyanide<br>0.1 M sodium cacodylate buffer, pH = 7.4 |
| DNA staining solution | 10 µM DRAQ5<br>0.1% SAPONIN<br>0.1 M sodium cacodylate buffer, pH = 7.4 |
| Bathing solution | 2.5 mM 3,3'- diaminobenzidine tetrahydrochloride (DAB)<br>0.1 M sodium cacodylate buffer, pH = 7.4 |
| Reduced osmium staining solution | 2% osmium tetroxide<br>1.5% potassium ferrocyanide<br>2 mM $CaCl_2$<br>0.15 M sodium cacodylate buffer, pH = 7.4 |
| Durcupan ™ resin mixture 1 | 10 mL Durcupan ™ ACM single component A, M, epoxy resin<br>10 mL Durcupan ™ ACM single component B, hardener 964<br>0.15 mL Durcupan ™ ACM single component D |
| Durcupan ™ resin mixture 2 | 10 mL Durcupan ™ ACM single component A, M, epoxy resin<br>10 mL Durcupan ™ ACM single component B, hardener 964<br>0.2 mL Durcupan ™ ACM, single component C, accelerator 960<br>0.15 mL Durcupan ™ ACM single component D |
| 1:1 infiltration mixture | 10 mL 100% ethanol<br>10 mL Durcupan ™ resin mixture 1 |
| 2:1 infiltration mixture | 5 mL 100% ethanol<br>10 mL Durcupan ™ resin mixture 1 |
